# Long-Term Outcomes of Pediatric Graves Disease Patients Treated with Anti-Thyroid Drugs: Experience from a Taiwan Medical Center

**DOI:** 10.1101/618942

**Authors:** Ya-Ting Chiang, Wei-Hsin Ting, Chi-Yu Huang, Shih-Kang Huang, Chon-In Chan, Bi-Wen Cheng, Chao-Hsu Lin, Yi-Lei Wu, Chen-Mei Hung, Hsin-Jung Li, Chia-Jung Chan, Yann-Jinn Lee

**Author notes:** Corresponding author (Lee YJ). These authors contributed equally to this work (Ya-Ting Chiang, Wei-Hsin Ting).

## Abstract

Graves disease (GD) is the most common cause of thyrotoxicosis in children and adolescents, accounting for 15% of all thyroid diseases during childhood. Anti-thyroid drugs (ATD) are recommended as the first-line treatment in children and adolescents. However, the remission rate is lower in children than in adults, and the optimal treatment duration and favorite factors associated with remission remain unknown. We aimed to investigate long-term outcomes of pediatric GD patients receiving ATD. We retrospectively reviewed medical charts of 300 pediatric GD subjects, who were initially treated with ATD and followed up for more than one year, from 1985 to 2017 at MacKay Children’s Hospital. The 300 patients comprised 257 (85.7%) females and 43 (14.3%) males, median age at diagnosis was 11.6 (range 2.7-17.8) years, and median follow-up period was 4.7 (range 1.1-23.9) years. Overall, 122 patients achieved the criteria for discontinuing ATD treatment, seventy-nine (39.9%) patients achieved remission, with a median follow-up period of 5.3 (range 1.5-20.1) years. Patients in the remission group were more likely to be aged < 5 years (remission vs. relapse vs. ongoing ATD; 11.4 vs. 0 vs. 2.6%, *P*=0.02), less likely to have a family history of thyroid disease (24.1 vs. 42.1 vs. 52.6 %, *P*=0.001), and had lower TRAb levels (42.8 vs. 53.6 vs. 65.1 %, *P*=0.02).

**Conclusion:** Long-term ATD remains an effective treatment option for GD in children and adolescents. Pediatric GD patients aged < 5 years, having no family history of thyroid disease and having lower TRAb levels were more likely to achieve remission.

## Introduction

Graves disease (GD) is a common disorder in adults, with a prevalence of approximately 0.5-1%. Pediatric patients account for < 5% of the total number of GD patients [1]. However, GD remains the most frequent cause of thyrotoxicosis in children and adolescents, accounting for 15% of thyroid disease during childhood [2]. The incidence increases gradually from young children and peaks in adolescents [3]. The optimal treatment option for GD in children and adolescents remains controversial. Current treatment approaches for GD include anti-thyroid drugs (ATD), radioactive iodine and surgery. ATD is usually recommended as the first-line treatment for GD in children and adolescents. However, the remission rate is lower in children than in adults [2, 4, 5]; the optimal treatment duration and the favorite factors associated with remission have not yet been established in children and adolescents [6–8].

The issue of how long ATD should be used in pediatric GD is important and warrants further study [5, 9]. In adult GD patients, if remission does not occur after 12-18 months of ATD therapy, the chance of remission with prolonged therapy is very low [4, 10]. In the pediatric population, longer treatment duration is associated with a higher remission rate. Lipple reported that the median time to remission with ATD was 4.3 years, and the expected remission rate was 25% every 2 years [11]. Leger reported that overall estimated remission rates after withdrawing ATD increased with time and were 20, 37, 45, and 49% after 4, 6, 8, and 10 years follow-up, respectively [6]. A retrospective study in Japan revealed that the remission rate was 46.2% after a median duration of 3.8 years [12]. However, long-term remission rate of pediatric GD cases treated with ATD was very low (< 20%), in cohorts from Australia [13] and Denmark [14].

Pediatric GD patients with some clinical or laboratory characteristics may have a higher chance of remission. A prospective study in France revealed that younger (age < 5 years), non-Caucasian children with severe initial presentation had a higher chance of relapse and required longer ATD treatment [7]. Another prospective study reported that initial less severe hyperthyroidism and the presence of other autoimmune conditions were remission predictors [6]. A similar study in the USA demonstrated that lower total T3, euthyroidism within 3 months of PTU and older age (age > 14.6 years) were significant remission predictors [8]; however, the largest retrospective study to date did not identify any significant factors to predict remission [12]. These above studies showed no consistent findings and few studies were conducted in the Asian population.

We aimed to investigate the long-term outcomes of pediatric GD patients who received ATD and identify probable clinical or laboratory factors associated with remission. We documented our 32-year experience in 300 children and adolescents with GD. Patients were classified as remission, relapse, and ongoing ATD groups; clinical and laboratory characteristics were presented and analyzed.

## Material and methods

We retrospectively reviewed medical charts of 396 GD subjects from 1985 to 2017 at MacKay Children’s Hospital. All the patients were diagnosed before 18 years of age. GD was diagnosed based on clinical and laboratory evidence, including thyrotoxicosis, diffuse goiter, with or without ophthalmopathy, elevated free T4/total T4 and suppressed TSH levels, and presence of autoantibodies against TSH receptor [4, 15]. Seventy-one patients were followed for less than one year, 6 patients received radioactive therapy and 19 patients received surgery as definite therapy and thus were excluded from our analyses. The remaining 300 patients initially treated with ATD and followed up for > 1 year constituted our study population.

We collected the following information from patients’ medical charts: age at diagnosis, sex, height, weight, body mass index (BMI), pubertal status, family history of thyroid disease (including autoimmune thyroid disease, goiter, thyroid nodule and thyroid cancer) in third-degree relatives, personal history of other autoimmune disease (type 1 diabetes, myasthenia gravis) or other syndrome (Down syndrome), initial free T4 (fT4), TSH receptor antibody (TRAb) levels, and interval until fT4, TRAb level normalized. All patients were initially treated with carbimazole or methimazole, with a starting dose between 2.5 and 30 mg/day, (0.05-0.80 mg/kg/day) depending on the patients’ age, body weight, clinical severity, and initial fT4 levels. PTU was only used when the patients could not tolerate the side effect of carbimazole or methimazole. The dose was subsequently titrated and adjusted to maintain euthyroidism. Patients were initially followed at 2-4 weeks interval and then every 3 months after thyroid function test results normalized. ATD was discontinued if euthyroidism was maintained at a low dose (methimazole ≤ 2.5 mg/day) for more than 6-12 months, and the TRAb was near or within the normal range. Remission was defined as the maintenance of euthyroidism ≥ 12 months after ATD was discontinued and no recurrence of thyrotoxicosis was recorded during the follow-up period. Relapse was defined as an elevated fT4, suppressed TSH levels together with restarting ATD use.

We obtained informed written consent from the parents or guardians of the children, and the study was approved by Mackay Memorial Hospital institutional review board (18MMHIS156e).

## Statistical analysis

We preformed descriptive statistics with categorical variables expressed as percentages and continuous variables as medians (25-75 percentiles) or means ± SD. *Univariate analysis.* A comparison of frequencies was performed employing the chi-square test or Fisher’s exact test (in case of expected frequencies < 5). A comparison of continuous variables was carried out using One-way ANNOVA or Kruskal-Wallis test while multiple groups were compared.

### Multivariate analysis

Multivariate logistic regression model was used to identify the possible remission predictors. Variables that were associated with remission in the univariate analysis and those judged to be potentially clinically relevant were entered the model. The variables used in the analysis were the proportion of young patients (age < 5 years), the proportion of patients with negative family history, initial fT4 levels, and TRAb levels at diagnosis. All the statistical analyses were performed using SAS software (version 9.4).

## Results

The 300 patients comprised 257 (85.7%) females and 43 (14.3%) males. Their median age at diagnosis was 11.6 (range 2.7-17.8), and 11 patients (3.7%) were diagnosed before the 5 years of age. The age and sex distributions were shown in Fig 1. One hundred and twelve patients (37.3%) reported a family history of thyroid disease. The median follow-up period of these patients was 4.7 (range 1.1-23.9) years. There were 102 patients (34%) who were lost follow-up during the study period. Those who were lost follow-up had no significant differences in the clinical and laboratory characteristics compared with those who remained in the study, except for shorter follow-up period (3.7 vs. 5.3 years, *P*=0.004).

**Fig 1.**
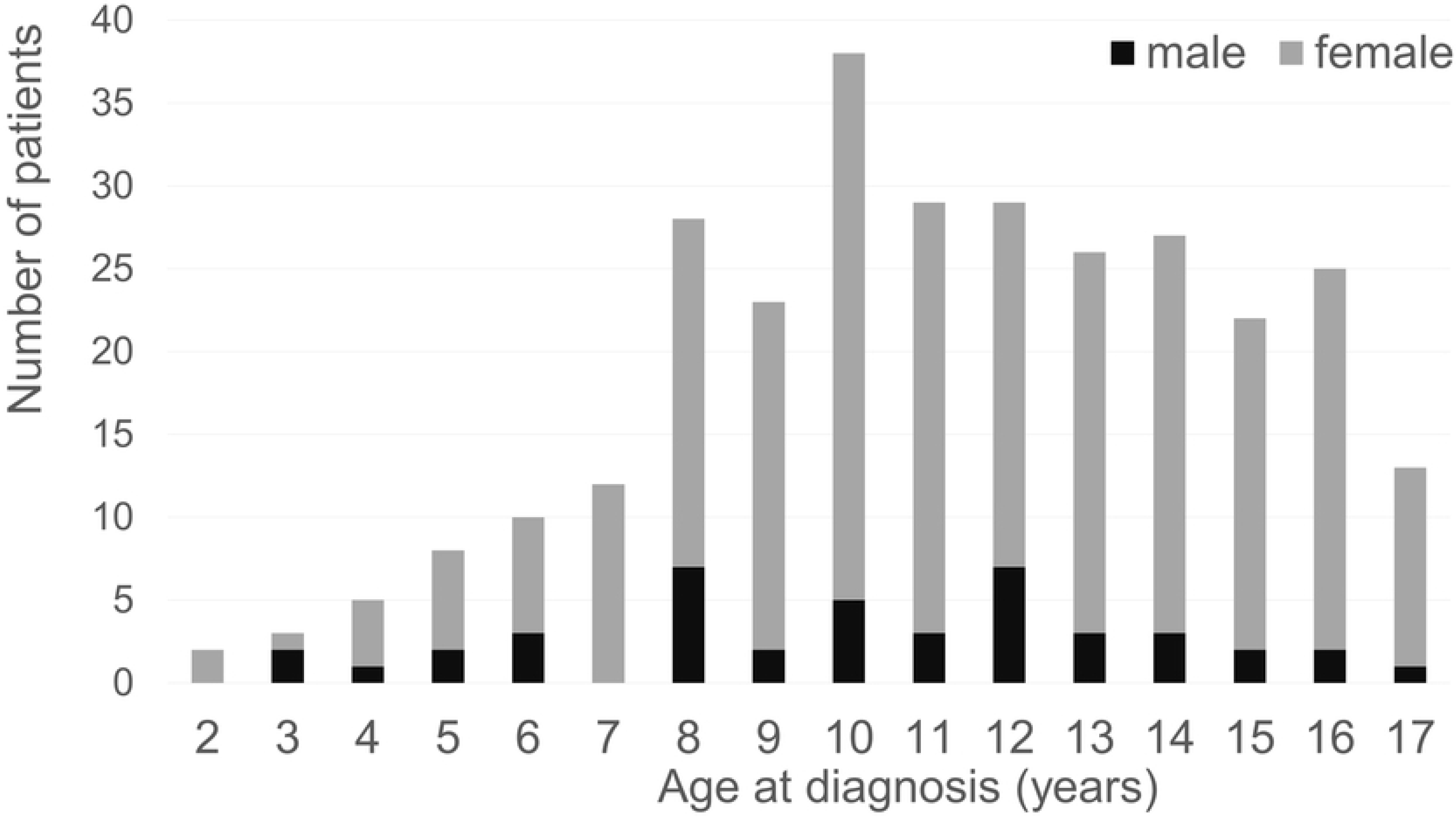
Age and sex distribution of children and adolescents with Graves disease (GD) diagnosis. The 300 patients consisted of 257 females and 43 males. The median age at diagnosis was 11.6 years (range 2.7–17.8 years). The incidence of GD increased markedly during adolescence.

From 198 patients who continued ATD treatment, ATD treatment was subsequently ongoing in 76 (38.4%) and was categorized as ongoing ATD treatment group. ATD was discontinued in 122 (61.6%) patients who met the criteria for discontinuing ATD treatment. Seventy-nine (39.9%) patients met the remission criteria, with a median follow-up of 5.3 (range 1.5-20.1) years and were classified as the remission group. Thirty-eight (19.2%) patients relapsed after ATD was discontinued, with a median of 0.7 (range 0.08-5.2) years and were assigned to the relapse group. The clinical course of the study population was shown in Fig 2.

**Fig 2.**
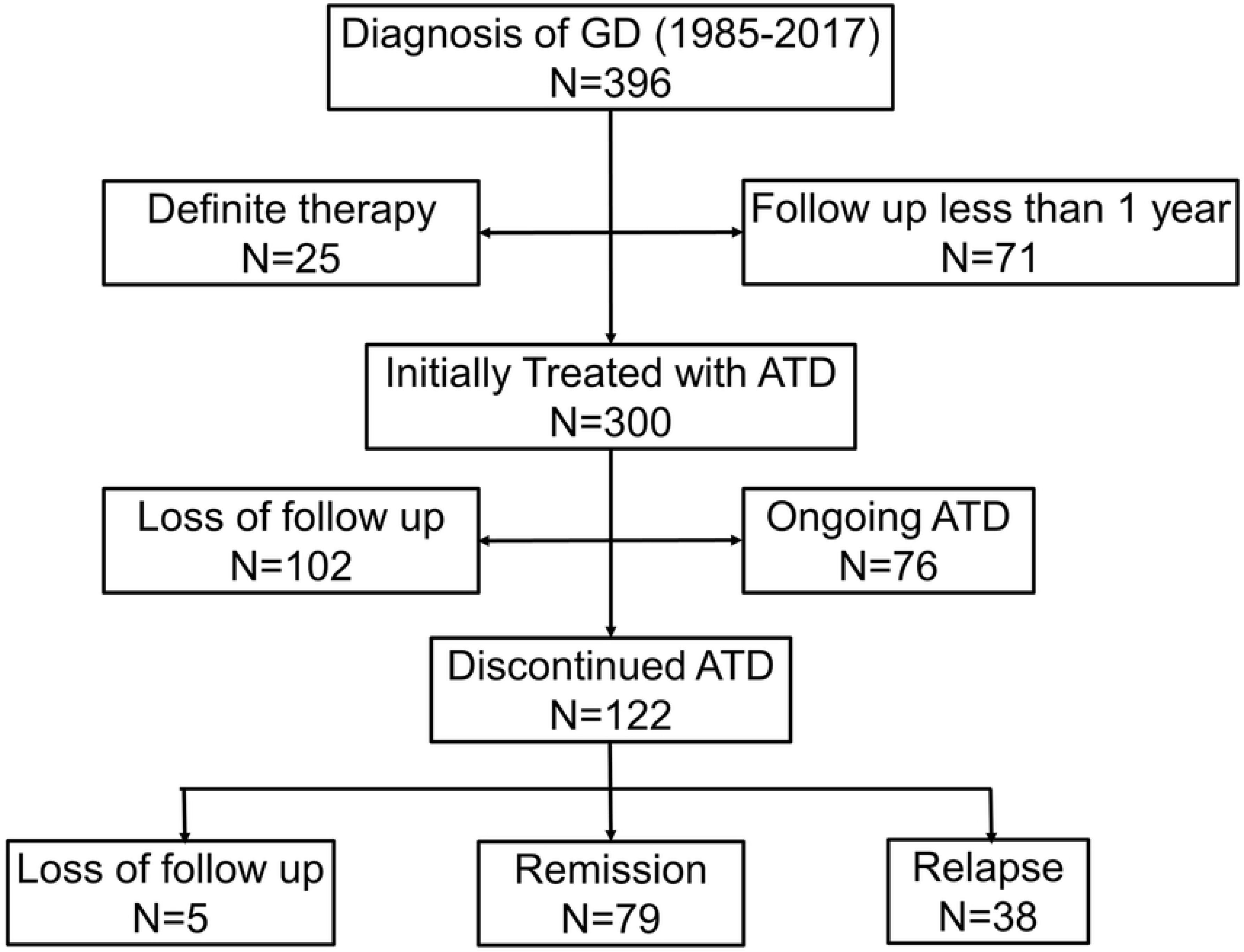
Clinical course of the study population initially treated with anti-thyroid drug (ATD). Of the 396 GD patients, 71 were followed for <1 year, 25 received definite therapy, and were excluded from our analyses. Of the remaining 300 patients, 102 patients were lost to follow-up during the study period. Of the 198 who continued ATD treatment, ATD treatment was subsequently ongoing in 76 (38.4%) and was discontinued in 122 (61.6%) patients who met the criteria of discontinuing ATD. Of the 122 patients who discontinued ATD treatment, 79 (39.9%) achieved a remission, 38 (19.2%) experienced a relapse, and 5 (2.5%) were lost to follow-up.

Patients in the remission group were more likely to be aged < 5 years (remission vs. relapse vs. ongoing ATD; 11.4 vs. 0 vs. 2.6%, *P*=0.02), less likely to have a family history of thyroid disease (24.1 vs. 42.1 vs. 52.6%, *P*=0.001), and had lower TRAb levels (42.8 vs. 53.6 vs. 65.1 %, *P*=0.02), (Table 1). In the remission group, patients aged < 5 years tended to receive ATD for a longer period than those with older age (younger vs. older age group: 7.2 vs. 5.0 years, *P*=0.28). The other variables, including male proportion, the proportion of puberty, height, weight and BMI z score, the proportion of patients with other diseases, initial ATD dose, initial serum fT4, and the interval until fT4 and TRAb levels became normal did not show any significant differences across the three groups.

**Table 1.**
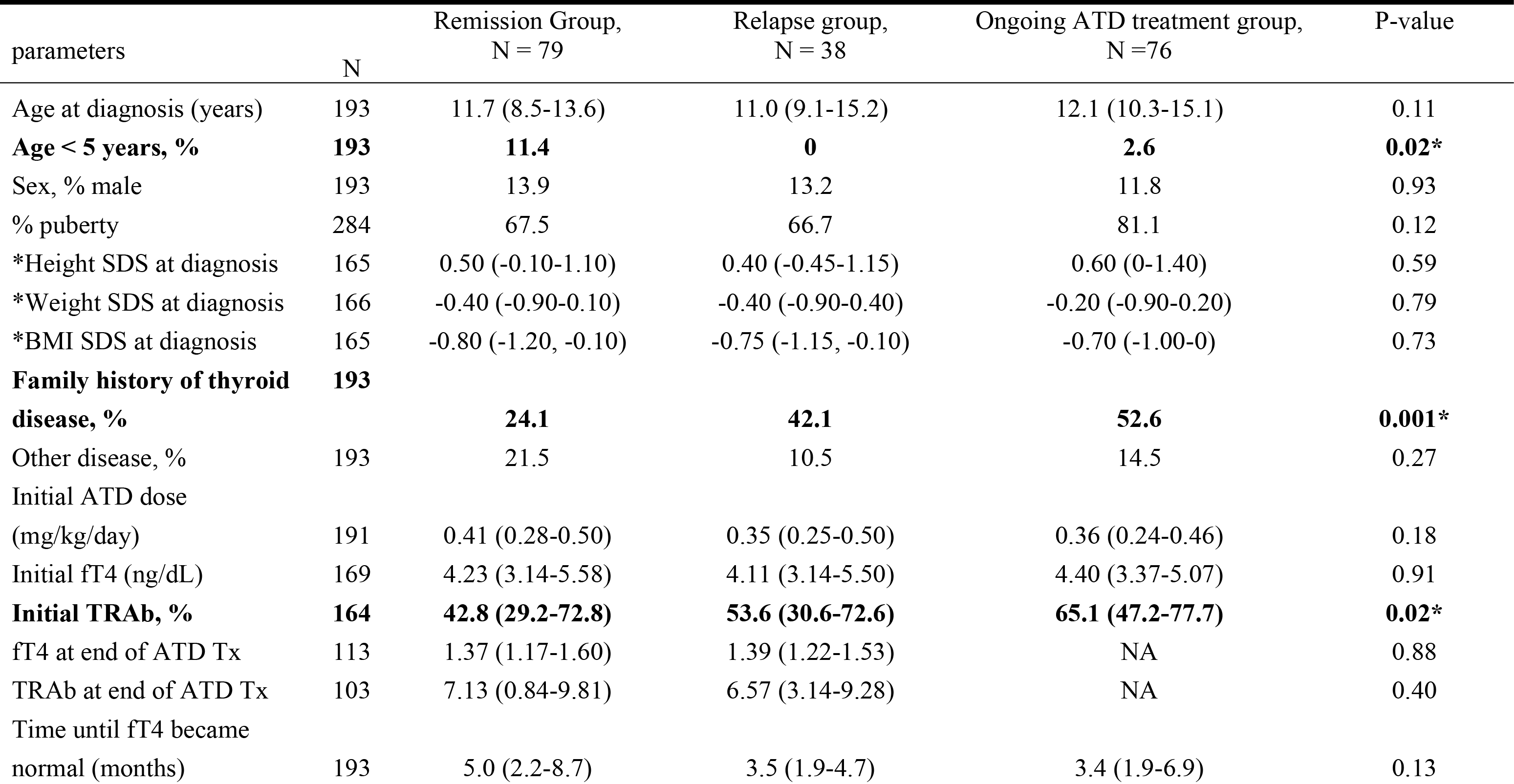
Clinical and Biochemical Characteristics of Four Groups: Remission Group, Relapse Group, and Ongoing Anti-Thyroid Drug Treatment Group

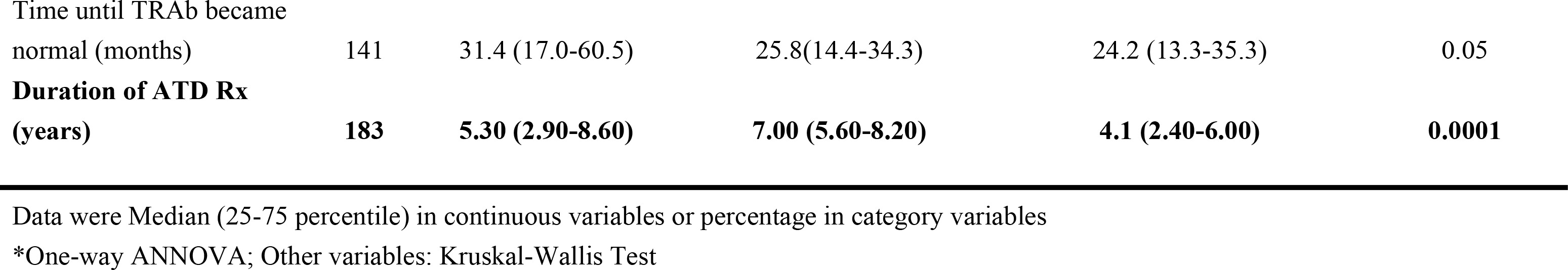

In the multivariate logistic regression model, we further identified patients who were aged < 5 years (Odds ratio [OR]: 12.6, 95% confidence interval [CI], 2.19-72.6; *P* = 0.005), had no family history of thyroid disease (OR: 3.75, 95% CI, 1.80-7.81; *P* = 0.0004), and had a lower initial TRAb levels (OR: 0.98, 95% CI, 0.97-0.99; *P* = 0.01) as remission predictors in pediatric GD (Table 2).

**Table 2.** Factors associated with remission in child and adolescents with Graves disease, multiple logistic regression model

| Variable | Univariate analysis |  | Multivariate analysis |  |
| --- | --- | --- | --- | --- |
|  | Odds ratio (95% CI) | P value | Odds ratio (95% CI) | P value |
| **Age < 5 years** | **7.20 (1.51-34.3)** | **0.01** | **12.6 (2.19-72.6)** | **0.005** |
| **No family history of thyroid disease** | **3.05 (1.62-5.74)** | **0.001** | **3.75 (1.80-7.81)** | **0.0004** |
| **Initial TRAb, %** | **0.98 (0.97-0.997)** | **0.01** | **0.98 (0.97-0.99)** | **0.01** |

## Discussion

In this study, we demonstrated that patients who aged < 5 years, who had no family history of thyroid disease and who had lower TRAb levels were more likely to achieve remission. There were 34% patients who were lost follow-up during the study period. Among patients who continued the ATD treatment, the long-term remission rate in pediatric GD patients was 39.9% after a median of 5.3 years of ATD treatment.

Our results suggested that GD patients who aged < 5 years had a higher chance to achieve remission and tended to receive ATD for a longer course. These young children were assumed to have better medical adherence under caregivers’ surveillance. Previous studies reported that pre-pubertal children needed a longer medical treatment and had a lower remission rate than pubertal children [16]. Lazar et al. reported that the remission rate was not different between pre-pubertal children and adolescents, but the time to remission tended to be longer in pre-pubertal children [17]. Two prospective studies showed that younger GD patients were less likely to achieve remission [8] or more likely to relapse after discontinuing ATD [7]. However, other retrospective studies performed in Japan [12] and Taiwan [18] did not determine age as a remission predictor. Because GD is rare in pre-pubertal children, especially in those aged < 5 years, further studies are needed to clarify the relationship between onset age and remission rate.

Our study showed that nearly 40% of pediatric GD patients had a family history of thyroid disease, consistent with previous studies in the literature [6, 7, 12, 19]. Our study also revealed that GD patients with a positive family history of thyroid disease were less likely to achieve remission. A similar study performed in 194 adult GD patients also proved that GD patients with a family history of thyroid disorders were 2.5 times more likely not to response to ATD treatment [20]. Although not all studies demonstrated a significant association between thyroid disease family history and the chance of remission [7, 8, 12, 21], our study still implied that family history acts as a remission indicator at the time of GD diagnosis.

Consistent with previous reports, our study also demonstrated that GD patients with lower titers of TRAb at diagnosis had a higher chance to remission [7, 21, 22]. TRAb was reported to be well correlated with GD severity and extra-thyroidal manifestations [23], showing concomitancy with the clinical course and being valuable for the diagnosis and management of children with GD [24]. A retrospective study conducted in 115 children aged 3-15 years showed that a TRAb level ≤ 2.5 times the upper reference limit, TRAb normalization during ATD and TRAb normalization time may predict further euthyroidism or hypothyroidism after ATD treatment stopped [21]. Pediatric GD patients with non-Caucasian origins, higher TRAb levels, higher free T4 levels, and younger age at diagnosis were reported to have a higher relapse rate [7]. These above studies, combined with our findings, confirmed TRAb as an indicator of GD activity and the predictive role for future remission occurrence after medical therapy.

Contradictory to adult GD cases, while a fixed course of ATD (no longer than 18 months) was recommended [25, 26], most studies showed that a longer ATD treatment duration increased remission rates in pediatric GD [2, 6, 7]. Recent published guidelines therefore suggested a prolonged course of ATD therapy before proceeding to definite therapy [26, 27]. However, the median time to remission reported in the literature was highly variable, and the optimal duration of ATD has not yet been determined. Our treatment protocol resulted in 40% of remission which is consisted with previous studies [12, 28], after a median of 5.3 years of ATD treatment. As shown in our study, long-term medical therapy resulted in high rates of lost follow up. Some clinicians believe that hypothyroidism is preferable to hyperthyroidism, because it is easier to treat and has a less serious morbidity [29]. However, medical adherence is problematic not only for long-term ATD therapy but also for the thyroxine supplement after hypothyroidism induced by definite therapy [30, 31]. Further long-term, prospective studies are required to determine the optimal duration of ATD treatment for pediatric GD.

There were several limitations in our study. The first limitation came from its retrospective nature, and high rates of lost follow up, which highlighted the difficulties in the daily practice. Since pediatric GD patients need a protracted ATD course to attain remission, meticulous and realistic counseling of patients and families should be started from the time of diagnosis [31]. Second, we did not analyze the patients’ characteristics who receiving radioactive iodine and total thyroidectomy, because few patients chose definite therapy in our institute, even in the relapse group. Third, the documentation of a family history of thyroid disease is not limited to autoimmune thyroid disease, which might introduce some bias to our estimate. Finally, the definition of remission is euthyroidism for only 12 months after ATD is discontinued. It is possible that patients experienced relapse one year after discontinuing medication. However, previous studies indicated that the risk of relapse declines with times [7, 12].

In conclusion, we identified pediatric GD patients who aged < 5 years, had no family history of thyroid disease and had lower TRAb levels were more likely to achieve remission. These remission predictors helped us to discuss with patients and families in the process of shared decision making and treatment plan. Long-term ATD is still a treatment option for pediatric GD, because our study showed that it resulted in a remission rate of 40%, with a median of 5.3 years ATD course. Such a long-term treatment course was inevitably associated with a poor medical adherence, realistic discussion and consultation should be applied in every newly diagnosed pediatric GD patients.

## Acknowledgments

This study was supported by grants RD1050151 from Mackay Medical College; and MMH 108-119 and MMH E-108-7 from MacKay Memorial Hospital, Taipei, Taiwan.

